# The limiting DNA replication initiation factors Sld2 and Sld3 influence origin efficiency independent of origin firing time

**DOI:** 10.1101/384644

**Authors:** Kelsey L. Lynch, Elizabeth X. Kwan, Gina M. Alvino, Bonita J. Brewer, M.K. Raghuraman

## Abstract

Chromosome replication in *Saccharomyces cerevisiae* is initiated from roughly 300 origins that are regulated both by DNA sequence and by the limited abundance of four *trans-acting* initiation proteins (Sld2, Sld3, Dpb11 and Dbf4, collectively called “SSDD”). We set out to determine how the association of Sld2 or Sld3 at origins contributes to time of origin activation and/or origin efficiency using auxin-induced protein degradation to further decrease their abundance. Depleting cells of either factor slows growth rate, increases S-phase duration, and causes viability defects, without activating the S phase checkpoint. Chr XII is uniquely unstable with breakage occurring specifically within the rDNA locus. The efficiency of the rDNA origin is decreased while the onset of replication initiation is unchanged. We found that origin efficiency is reduced uniformly across the unique portions of the yeast genome. We conclude that the abundance of Sld2 and Sld3 contribute primarily to origin efficiency.

## Introduction

One of the enduring mysteries in the field of eukaryotic DNA replication is the variability in onset, or initiation, of DNA replication across a genome. Eukaryotes initiate DNA replication from multiple sites, known as origins, which are distributed along the length of each linear chromosome (Cairns, 1966; reviewed by Prioleau and MacAlpine, 2016). Early studies of DNA replication revealed that certain parts of the eukaryotic genome are replicated earlier in S phase than others, and that the large-scale organization of chromosomes, their underlying sequence and chromatin structure play a regulatory role in this variability (Lima-de-Faria and Jaworksa, 1968; Taylor, 1960). These differences in replication time arise from staggered origin firing during S phase with initiation occurring in a continuum from early to later in S phase (Ferguson et al., 1991; Raghuraman et al., 2001). Furthermore, not all potential sites of initiation are used in every cell and some are completely inactive in a native chromosomal context (Brewer and Fangman, 1988; Newlon et al., 1991)—a characteristic referred to as origin “efficiency”. In broad terms, these two modes of variability in replication initiation, origin efficiency and origin firing time, are conserved from *Saccharomyces cerevisiae* to humans (Koren et al., 2014; Raghuraman et al., 2001). The conservation of these features implies that plasticity in origin activation is biologically relevant, although the nature of its importance for genome function is not entirely clear.

The proteins that recognize origin DNA and initiate replication at those loci are well conserved across eukaryotes; however, the molecular details have been most comprehensively studied in the budding yeast *S. cerevisiae* (reviewed by Bell and Labib, 2016). For an origin to fire, the origin DNA sequence is first bound by the origin recognition complex (ORC) which, through a set of interacting protein factors, coordinates the loading of the replicative MCM2-7 helicase during G1 (Bell and Stillman, 1992; Diffley and Cocker, 1992; Diffley et al., 1994; Sun et al., 2013). In the transition from G1 to S phase, a series of initiation factors, a subset of which are illustrated in Figure 1A, associate transiently with the assembling complex resulting in the assembly of the active replicative helicase known the CMG (Cdc45-MCM2-7-GINS) helicase (Douglas et al., 2018; Ilves et al., 2010). CMG assembly is considered the commitment step of replication initiation at a single origin and leads irreversibly to the establishment of bidirectional replisomes (Bell and Labib, 2016). Variation in this crucial replisome assembly step determines time of origin firing and origin efficiency.

**Figure 1.**
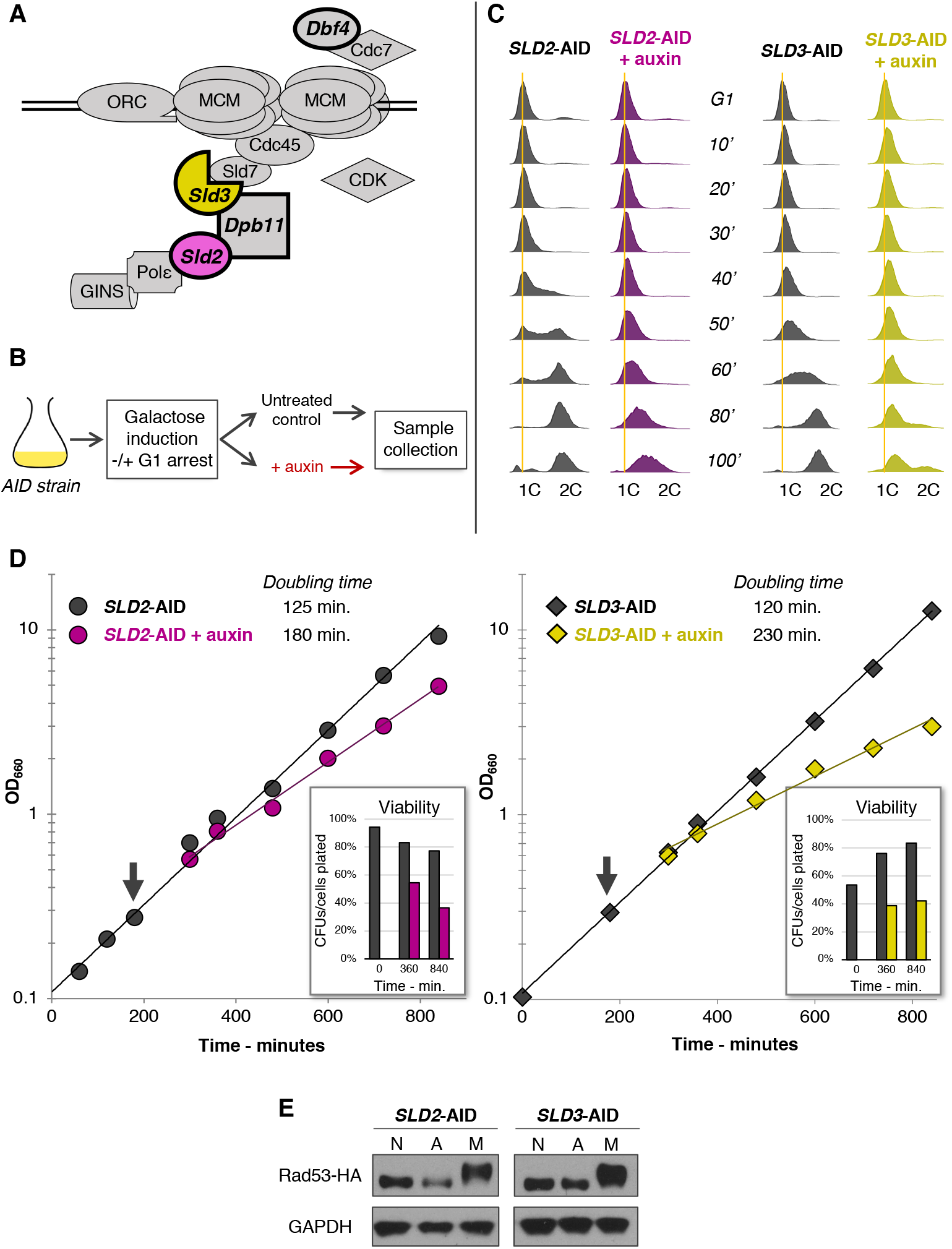
Initial phenotyping of Sld2- or Sld3-depleted cells. (A) Sld2 and Sld3, shown in purple and yellow, are two replication initiation factors thought to limit origin firing in *S. cerevisiae*. Here, they are shown with the assembling replisome. Two other limiting initiation factors, Dbf4 and Dpb11, are shown in bold. (B) General outline for experiments using the AID system to degrade a protein of interest. (C) Flow cytometry profiles for synchronized S phase in *SLD2*-AID and *SLD3*-AID. The vertical orange lines indicate 1C DNA. (D) Log phase growth of *SLD2*-AID and *SLD3*-AID. The arrow indicates when we divided the cultures and added auxin. Sld2 depletion increased the doubling time by 44%. Sld3 depletion increased the doubling time by 92%. The proportions of cells able to form colonies, i.e. the proportion of viable cells after being returned to no-depletion conditions, are shown for three timed samples. (E) Western blot for Rad53 phosphorylation in *SLD2*-AID and *SLD3*-AID strains containing an HA-tagged version of Rad53. Samples are untreated, asynchronously growing cells (lanes N), treated with 0.5 mM auxin for 2 hours (lanes A), or treated with 0.1% MMS for 1 hour as a positive control (lanes M).

The assembly and activation of replisomes are regulated by two kinases, S-CDK (S-phase cyclin-dependent kinase; Cdc28 complexed with either Clb5 or Clb6) and DDK (Dbf4-dependent kinase; Cdc7 complexed with Dbf4) (Bousset and Diffley, 1998; Donaldson et al., 1998; Enserink and Kolodner, 2010; Jackson et al., 1993; Labib, 2010). Clb5 and Clb6 have redundant roles in phosphorylating various replisome components; however, since Clb6 is expressed only at the beginning of S phase, late origin firing is dependent on Clb5 expression (reviewed by Bloom and Cross, 2007; Donaldson et al., 1998; McCune et al., 2008).

The processes that distribute CDK and DDK to specific origins are not well understood. The underlying DNA sequences and chromatin context at origins seem to have the greatest influence (Lima-de-Faria and Jaworksa, 1968; Pohl et al., 2012; reviewed by Rhind and Gilbert, 2013; Vogelauer et al., 2002). However, the discovery that four specific replication initiation factors (Sld2, Sld3, Dpb11, and Dbf4—the “SSDD” factors) are produced at concentrations significantly lower than the rest of the origin/replisome components suggested that they may be rate-limiting for origin initiation (Mantiero et al., 2011). When Mantiero et al. (2011) overexpressed all four proteins simultaneously, the temporal firing of origins was compressed, and a subsequent study revealed that origin efficiency increased genome-wide (McGuffee et al., 2013). These findings suggest that the varying ability of different parts of the genome to efficiently recruit the SSDD factors results in the pattern of origin use observed during DNA replication. From their observations, Mantiero et al. (2011) developed a model whereby genome features near origins determine how well origins are able to recruit the limiting SSDD factors, with the earliest and most efficient origins being the most competitive for recruiting SSDD factors. Since each of these four proteins is required only for the initiation phase of DNA replication (Bell and Labib, 2016; Deegan et al., 2016; Muramatsu et al., 2010; Tanaka et al., 2007; Zegerman and Diffley, 2007), Mantiero et al. (2011) proposed that as early and/or efficient origins become active and release the limiting factors from the assembling replisome, those factors are recycled to origins that are less competitive for recruiting SSDD and thereby promote origin firing at later and/or more inefficient origins.

More recently, Collart et al. (2013) demonstrated that the same SSDD factors restrict origin firing during early embryonic development in *Xenopus laevis*. At the mid-blastula transition (MBT), developing embryos transition from relying on stores of maternally-derived transcripts to activating embryonically-driven transcription (Farrell and O’Farrell, 2014; Langley et al., 2014). The transcriptional activation of the genome results in drastic changes in embryonic cell cycles—from rapid, synchronous divisions with fast S phases and an almost complete lack of gap phases to slower, asynchronous divisions with G1 and G2 phases and a slowed S phase featuring greatly reduced origin activity. Collart et al. (2013) demonstrated that the reduction in DNA replication initiation at the MBT in *Xenopus* depends upon titration of the orthologous SSDD factors. By simultaneously increasing the concentrations of the SSDD factors, they found that replication initiation was not reduced, that the cells continued to divide rapidly, and that the embryos failed to develop. These observations suggest that the establishment of variation in origin activity is an essential feature of embryogenesis.

While the SSDD factor model for variable origin firing time and efficiency is an attractive one, there are still many questions regarding the mechanisms of how the SSDD factors associate with origins. Do some origins have a more favorable underlying DNA sequence, a more favorable chromatin conformation, and/or reside in a prime nuclear compartment that promotes their advanced time of firing or their efficiency? Do some of the SSDD factors contribute to efficiency while others contribute to the temporal aspect of origin firing?

For our study, we have explored the role that SSDD factors have in determining origin efficiency and firing time. One feature that influenced our work was the observation that even when the SSDD proteins were overexpressed in yeast, there were still differences in time of replication among the origins assayed (Mantiero et al., 2011). This observation, coupled with the knowledge that each of the SSDD factors interacts with the replisome at different times and with different components of the complex (Bell and Labib, 2016; Kamimura et al., 2001; Muramatsu et al., 2010; Sheu and Stillman, 2010; Zegerman and Diffley, 2007), suggested that the abundance of these factors may contribute differently to origin firing time and efficiency. Crucially, all four SSDD factors had to be expressed simultaneously to advance replication kinetics (Mantiero et al., 2011), indicating that the abundance of any single factor could limit the number of active origins in a cell. Therefore, we reasoned that reducing the abundance of any one SSDD factor would restrict origin firing below WT levels.

To test our prediction, we induced degradation of individual SSDD factors (Sld2 or Sld3) then assessed changes in cell cycle progression, genome instability, and chromosome replication. We find that reduced abundance of either Sld2 or Sld3 results in viability defects, and that Chromosome XII is particularly unstable as a consequence of the failure to complete rDNA replication. Significantly, even though initiation of the rDNA origin is impaired, the rDNA origins still can fire at their normal time in S phase. Likewise, we find that origin firing is depressed across the genome, but we do not observe any changes in origin firing time. We conclude that Sld2 and Sld3 abundance both impact origin efficiency rather than time of replication initiation. Additionally, we believe that our work has implications for the ongoing debate in the DNA replication field surrounding a recently developed model for replication initiation termed the “stochastic model for origin firing.”

## Results

### Independent depletion of two of the SSDD limiting replication initiation factors slows S phase progression, reduces growth rate, and impacts cellular viability

In this study, our goal was to reduce the abundance of individual SSDD factors, one at a time, and then examine DNA replication initiation genome-wide to test whether those factors contribute to time and/or efficiency of origin activity. Using the 3x-mini-AID version of the auxin inducible degron (AID) system (Kubota et al., 2013; Nishimura et al., 2009), we set out to engineer strains with a C-terminal auxin-inducible degron tag on either of two SSDD factors, Sld2 and Sld3, referred to as *SLD2*-AID and *SLD3*-AID. Because Sld2 and Sld3 associate sequentially with the replisome during the cell cycle and interact with distinct proteins within the replication complex (Figure 1A) (reviewed by Bell and Labib, 2016; Kamimura et al., 2001; Tanaka et al., 2011), we expected that focusing on those factors could reveal insights into how the SSDD proteins contribute uniquely to variable replication initiation. Note that henceforth, by “induced” *SLD2*-AID or *SLD3*-AID we mean depletion of Sld2 or Sld3, respectively, while “uninduced” indicates the control, no-depletion condition.

We tested S-phase progression in *SLD2*-AID and *SLD3*-AID by measuring DNA content with flow cytometry. After arresting cells in G1 phase and inducing expression of the plant-derived E3 ubiquitin ligase with galactose, we began degradation of Sld2 or Sld3 by adding 0.5 mM auxin to one half of the culture and leaving the other half as an uninduced control (Figure 1B). Both uninduced *SLD2*-AID and *SLD3*-AID completed S phase by 80 minutes, with *SLD3*-AID entering S phase more slowly than the *SLD2*-AID strain (Figure 1C). The S phase delay in uninduced *SLD3*-AID may be the consequence of the degron tag interfering with Sld3 function. However, the presence of auxin severely impeded progress through S phase for both strains: by 100 minutes neither strain had reached 2C DNA content, again with *SLD3*-AID lagging behind *SLD2*-AID (Figure 1C). We suspect that the more pronounced S phase delay resulting from Sld3 depletion compared to the delay caused by Sld2 depletion may reflect a key difference between the two proteins. Sld3 is less abundant than Sld2, even before depletion, so reducing its abundance further may suppress origin firing more severely (Ghaemmaghami et al., 2003; Mantiero et al., 2011). To rule out possible off-target effects of the plant-derived E3 ubiquitin ligase expression, we carried out the same experiment in a strain harboring the *GAL1-10*-*Os*TIR1-9myc construct alone, without a degron domain-tagged target protein (referred to as the “AID parent strain”). We observed slight differences in cell cycle progression after adding auxin to the AID parent strain, but there were no discernible changes in S phase progression (Supplemental Figure 1A).

To determine whether the increased S phase duration we observed in the Sld2- and Sld3-limited strains impacts growth rate and/or cellular viability, we compared log-phase growth of the two degron strains. Doubling times increased for both *SLD2*-AID and *SLD3*-AID when the target proteins were degraded (Figure 1D). In agreement with the more severe S phase phenotype observed by flow cytometry, the Sld3-depleted cells exhibited a greater increase in doubling time compared to cells depleted for Sld2 (Figure 1D). In the final timed sample, both strains had experienced an approximately 40% decrease in viability compared to the uninduced control as assayed by colony-forming ability of cells (Figure 1D). While auxin treatment did slightly slow growth in the AID parent strain, the difference was not due to decreased viability (Supplemental Figure 1B).

One explanation for the rapid loss of viability in the auxin-induced cultures is that the S phase checkpoint is activated in response to initiation factor depletion, preventing the cells from completing the cell cycle and eventually leading to cell death (Foiani et al., 2000; Zhang and Hunter, 2014). To test for checkpoint activation, we HA-epitope tagged Rad53 in the AID strains, then tested for phosphorylation of Rad53 during asynchronous growth. After two hours in auxin, *SLD2*-AID and *SLD3*-AID both had S phase defects, but the AID parent did not (Supplemental Figure 1C). We conclude that the stalled S phase was not caused by S phase checkpoint activation because we did not observe Rad53 phosphorylation in the auxin-treated samples of either strain (Figure 1E). Likewise, adding auxin to the AID parent did not lead to Rad53 phosphorylation (Supplemental Figure 1D).

### Chromosome XII is specifically destabilized by Sld2 or Sld3 depletion

Although the S phase checkpoint is not active when Sld2 or Sld3 are depleted, the rapid loss of viability suggested that genome stability may nonetheless be compromised. Successful activation of the S phase checkpoint requires that cells establish a threshold of active replication forks (Shimada et al., 2002; Tercero et al., 2003). We hypothesized that Sld2 or Sld3 depletion might reduce origin firing below the threshold required for S phase check point activation and, as a result, some cells would enter mitosis with partially-replicated chromosomes causing double-stranded breaks, mitotic catastrophe, and cell death (Feng et al., 2011).

We used counter-clamped homogeneous electric field (CHEF) electrophoresis to assess chromosome stability during and after a synchronous S phase in *SLD2*-AID and *SLD3*-AID. Branched chromosomes formed during replication are unable to migrate from the wells and, therefore, the hybridization signal for individual chromosomes at their characteristic positions on CHEF gels is reduced while signal in the well is increased (Cha and Kleckner, 2002; Hennessy et al., 1991). If chromosome replication is slowed, we would expect increased hybridization at the well and for well hybridization to be present for a longer period of time. If partially-replicated chromosomes break from mitotic catastrophe, we would expect diffuse hybridization signal below the position of the chromosome band.

We synchronized *SLD2*-AID and *SLD3*-AID cultures in G1 and released them into S phase with and without auxin. We monitored cell cycle progression by flow cytometry and collected samples at intervals for 420 minutes (Figure 2A). Following the 420-minute sample collection, we allowed the remainder of the culture to grow to stationary phase overnight before recovering the final sample. We performed the same experiment using the AID parent strain (Supplemental Figure 2C, D). All timed samples were embedded in agarose plugs and chromosomes examined by CHEF gel electrophoresis.

**Figure 2.**
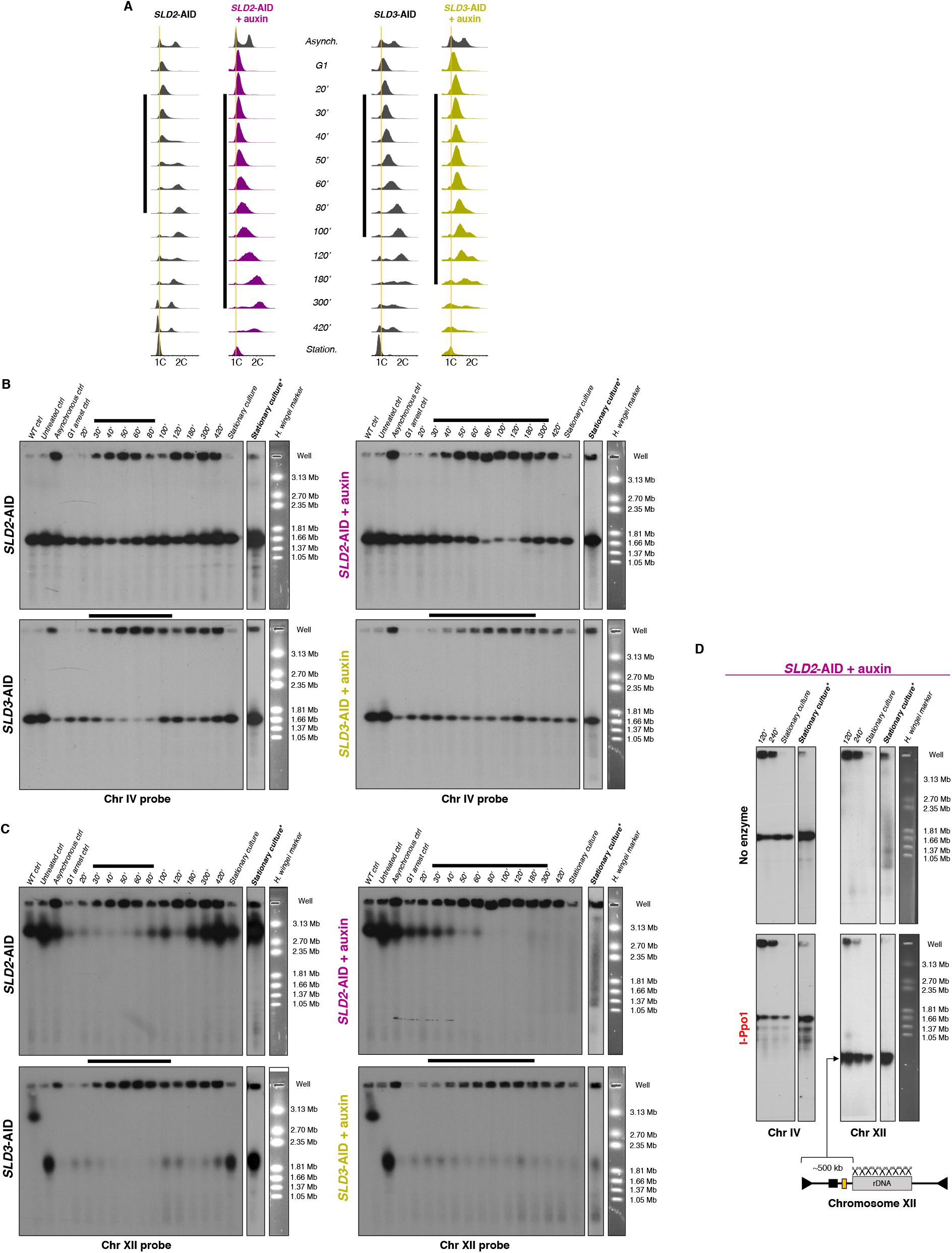
CHEF gel analysis of chromosome stability during Sld2- or Sld3-depleted S phase. (A) Flow cytometry profiles for the synchronized *SLD2*-AID and *SLD3*-AID strains, as well as asynchronous and stationary phase cells, collected for CHEF gel analysis. The vertical black bar to the left of each set of profiles shows the estimated duration of synchronous S phase. We continued to collect samples until synchrony was lost after T100 in the uninduced cells. Note that in the auxin-treated samples there is a small peak at 1C DNA in the later samples suggesting that the cells eventually divided. (B) Southern blot of the CHEF gels probed for Chr IV (*GAL3* probe). The horizontal black bars correspond with the synchronous S phase samples shown in (A). The EtBr-stained gel images to the right show *Hansenula wingei* chromosomes (sizes in Mb) as markers for electrophoretic mobility. Hybridization at the top of each lane is from DNA trapped in the wells. The “WT control” is a stationary phase W303 sample. The “untreated control” is from stationary phase cells that were neither galactose induced, nor auxin treated. The “asynchronous control” is from cells before a factor arrest. “Saturated culture” is from cells that grew past 420 minutes to stationary phase. The images of the “saturated culture” sample indicated by bold type and an asterisk are longer film exposures to reveal subchromosome-sized DNA fragments. (C) The blots in panel B were re-probed with a single copy sequence on Chr XII (*CDC45*). (D) Chr IV- and Chr XII-probed Southern blots of auxin-induced, asynchronous *SLD2*-AID strain samples with and without I-Ppo1 restriction enzyme treatment. The illustration of Chr XII below the blots indicates where I-Ppo1 cuts within the rDNA on Chr XII and the predicted size for the CDC45-containing fragment to the left of the rDNA. The yellow rectangle shows the approximate location of *CDC45*.

We probed the Southern blots of the CHEF gels for single-copy sequences on individual chromosomes (Chr IV in Figure 2B; blot images and quantification of chromosomes XV, X and III in Supplemental Figures 2G – J). We prepared the WT control and untreated control samples from stationary phase (non-cycling) cells and thus there was little hybridization to the wells (Figure 2B). In the asynchronous samples, we observed more Chr IV signal in the well, reflecting the higher proportion of S phase cells (Figure 2B). The fraction of Chr IV signal migrating at its normal location was reduced as cells entered S phase and the well signal increased (Figure 2B). By T80 - T100, we found the majority of signal returned to the expected Chr IV position and signal at the well was reduced (Figure 2B). The period between T30 and T80 marks the duration of replication for Chr IV (Figure 2B) and corresponds well with the estimated S phase length we determined by flow cytometry (Fig 2A, left). We quantified the well signal as a proportion of total chromosome signal to highlight the changes in well-retention over the course of S phase (Supplemental Figure 2A and 2B). At the latest times (T180 – T420), when the uninduced cells had lost synchrony but were still in log-phase growth, the ratio of signal from partially-replicated and fully-replicated chromosomes was constant (Figure 2B and Supplemental Figure 2B).

In response to depletion of Sld2 or Sld3, the distributions of hybridization signal in the well vs. the signal from full-length Chr IV (Figure 2B; Supplemental Figure 2B) were in agreement with the flow cytometry analysis, indicating that Chr IV takes longer to finish replication when Sld2 or Sld3 are depleted. The accumulation of Chr IV signal in the wells was delayed (T30 – T180 in Figure 2B and Supplemental Figure 2B), and the maximum well signal occurred between T80 – T100, compared to T50 – T60 in the control (Figure 2B and Supplemental Figure 2B). We did not see evidence of random breakage for Chr IV in the timed or the stationary phase samples even after long exposures of the blot (Figure 2B). Additionally, we saw no differences in Chr IV migration depending on the presence of auxin in the AID parent strain control (Supplemental Figure 2C, D). We conclude that Chr IV replication takes proportionately longer when either Sld2 or Sld3 are depleted, but replication of that chromosome appears to be complete since there is no evidence of breakage as the cells enter mitosis. Thus, Chr IV is not destabilized by depletion of either Sld2 or Sld3. The same conclusions can be made from our analyses of chromosomes XV, X, and III (Supplemental Figure 2G - J). In contrast, the pattern of replication and stability of chromosome XII were strikingly different.

In uninduced *SLD2*-AID, the migration pattern of Chr XII was similar to that of Chr IV, although there was generally more Chr XII signal trapped in the well in all timed samples (Figure 2C, top left, and Supplemental Figure 2E). Increased well-retention of Chr XII compared to other chromosomes is thought to be due in part to its greater length (~2X times the length of Chr IV) and to unresolved branched recombination intermediates within the rDNA locus (Ide et al., 2007). As cells entered S phase, the majority of Chr XII signal remained in the well and returned to its full-length position at the end of S phase in agreement with the S phase kinetics (Figure 2A, left). However, upon depletion of Sld2, Chr XII became trapped in the well and did not return to its full-length position in the gel (Figure 2C). By the time the culture reached stationary phase, roughly half of the Chr XII signal was in subchromosomal-sized fragments (Figure 2C). It appears that Chr XII, in response to Sld2 depletion, never completed replication and, as cells entered mitosis, was fragmented.

*SLD3*-AID cells had similar problems maintaining Chr XII. First, we noted that Chr XII in the *SLD3*-AID strain is more than 1.5 Mb shorter that the parental wild type strain (Figure 2C, left bottom). Because *SLD3*-AID is haploid, the reduction in size can only be explained by a loss of rDNA repeats. We estimate that there are ~250 rDNA repeats in the *SLD2*-AID strain vs. ~90 copies in the *SLD3*-AID strain. Consistent with the duration of S phase (Figure 2A), Chr XII signal persisted in the well longer in untreated *SLD3*-AID relative to *SLD2*-AID, potentially reflecting compromised function of degron-tagged Sld3 (Figure 2C; Supplemental Figure 2E). But by T100 in uninduced *SLD3*-AID, full length Chr XII returned to its position in the gel (Figure 2C). Post Sld3 depletion, most of the Chr XII signal remained in the well and appeared as subchromosomal fragments in the stationary phase culture (Figure 2C). The presence of small amounts of full-length Chr XII in the later samples (T100 – T300 in Figure 2C and Supplemental Figure 2E) likely originated from the subpopulation of cells remaining in G1 (Figure 2A, far right).

The persistence of Chr XII signal in the well followed by the appearance of subchromosomal fragments suggests that only Chr XII is unable to complete replication before the onset of anaphase resulting in chromosome-specific breakage. Our additional finding that, even without auxin treatment, *SLD3*-AID is unable to maintain the normal ~250 copies of the rDNA repeat suggests that the problem resides specifically with rDNA replication. If the problem were specific to the rDNA and not widespread across Chr XII, we would expect that the branched DNA that prevents Chr XII from migrating in the gel would be present only within the rDNA itself and not in the flanking portions of the chromosome.

To test the possibility that branched intermediates were restricted to the rDNA, we digested agarose-embedded DNA with I-Ppo1—a restriction enzyme that claves in each rDNA repeat but nowhere else in the yeast genome (Ellison and Vogt, 1993; Promega, 2018)—and then determined whether the unique portions of Chr XII were able to enter the gel. If incomplete replication of the rDNA causes persistent branched structures on Chr XII, then cutting the chromosome within the rDNA should release the fully-replicated regions on either side of the rDNA and allow those arms of the chromosome to migrate into the gel. Probing the Southern blot for sequences outside of the rDNA would reveal whether the non-rDNA portions of Chr XII are able to migrate after the rDNA repeats are digested. Accordingly, we repeated the CHEF gel collection, this time using asynchronous cultures of the *SLD2*-AID strain and collecting cells 120 minutes, 240 minutes after adding auxin, and after growth to saturation. Flow cytometry confirmed an enrichment of S phase cells in the 120- and 240-minute samples (Supplemental Figure 3).

I-Ppo1 treatment of the 120- and 240-minute samples restored migration of the leftmost portion of Chr XII (Figure 2D). In the stationary phase sample, the Chr XII region left of the rDNA migrated as a discrete band of the expected size after I-Ppo1 digestion, suggesting that chromosome breakage occurred specifically in the rDNA (Figure 2D). The retention of Chr XII in the CHEF gel wells and the release of the hybridization signal by I-PpoI cleavage suggest that the specific fragmentation of Chr XII and the reduction in viability are the consequence of incomplete replication of the rDNA when Sld2 or Sld3 are further limited.

### Depletion of Sld2 or Sld3 reduces origin efficiency of the high copy rDNA origin of replication without delaying the onset of initiation

Since Sld2 and Sld3 are both required for assembly of the replisome at origins, we reasoned that the rDNA replication problem might occur during initiation—either reduced efficiency of origin activation or delayed activation, such that replication of the locus cannot be completed in a timely manner. Each rDNA repeat contains a potential origin of replication within the non-transcribed spacer that lies between the 5’ ends of the divergently transcribed 35S and 5S genes (reviewed by Venema and Tollervey, 1999). On a recombinant plasmid, the rDNA origin is inherently inefficient, a property that is attributed to a single nucleotide replacement within the origin consensus sequence (Kouprina and Larionov, 1983; Kwan et al., 2013; Miller and Kowalski, 1993). The combination of this SNP, silencing of the endogenous rDNA locus by Sir2 and other chromatin features unique to the rDNA, and the variable 35S transcriptional activity found across the rDNA array is thought to limit the origin firing density of the rDNA in its native context to about one origin in every 3-5 repeats (Brewer and Fangman, 1988; Fritze et al., 1997; Muller et al., 2000; Yoshida et al., 2014).

We assessed rDNA origin activity by examining the rDNA repeats, collapsed as a single restriction fragment using 2D gel electrophoresis. The replication intermediates from the collection of identical restriction fragments are branched molecules that contain either a bubble shape (indicating initiation) or a Y shape (indicating passive replication). We collected samples from asynchronous, G1-arrested, and S phase-timed samples for uninduced and induced *SLD2*-AID and *SLD3*-AID cells then analyzed replication intermediates by 2D gel for the NheI restriction fragment containing the rDNA origin (Figure 3A and 3B, respectively). A decrease in efficiency at the rDNA origin would cause a reduction in replication bubble signal relative to replication fork signal on the Southern blot of the 2D gel, whereas a delay in activation would be manifested as a difference in the time at which bubble structures first become apparent when compared to the control condition (Figure 3). The uninduced cells show initiation beginning by T30 and falling to nearly undetectable levels by T90, which is when cells are at or near 2C DNA content. We quantified the bubble:1N ratio for each timed sample (Figure 3C, D, E). The signal from replication bubbles from uninduced *SLD2*-AID peaked at T30 and gradually diminished as the cells neared 2C DNA content (Figure 3D). For uninduced *SLD3*-AID, the bubble signal peaked later (T50 - T60) and declined to barely detectable levels by T90 (Figure 3E).

**Figure 3.**
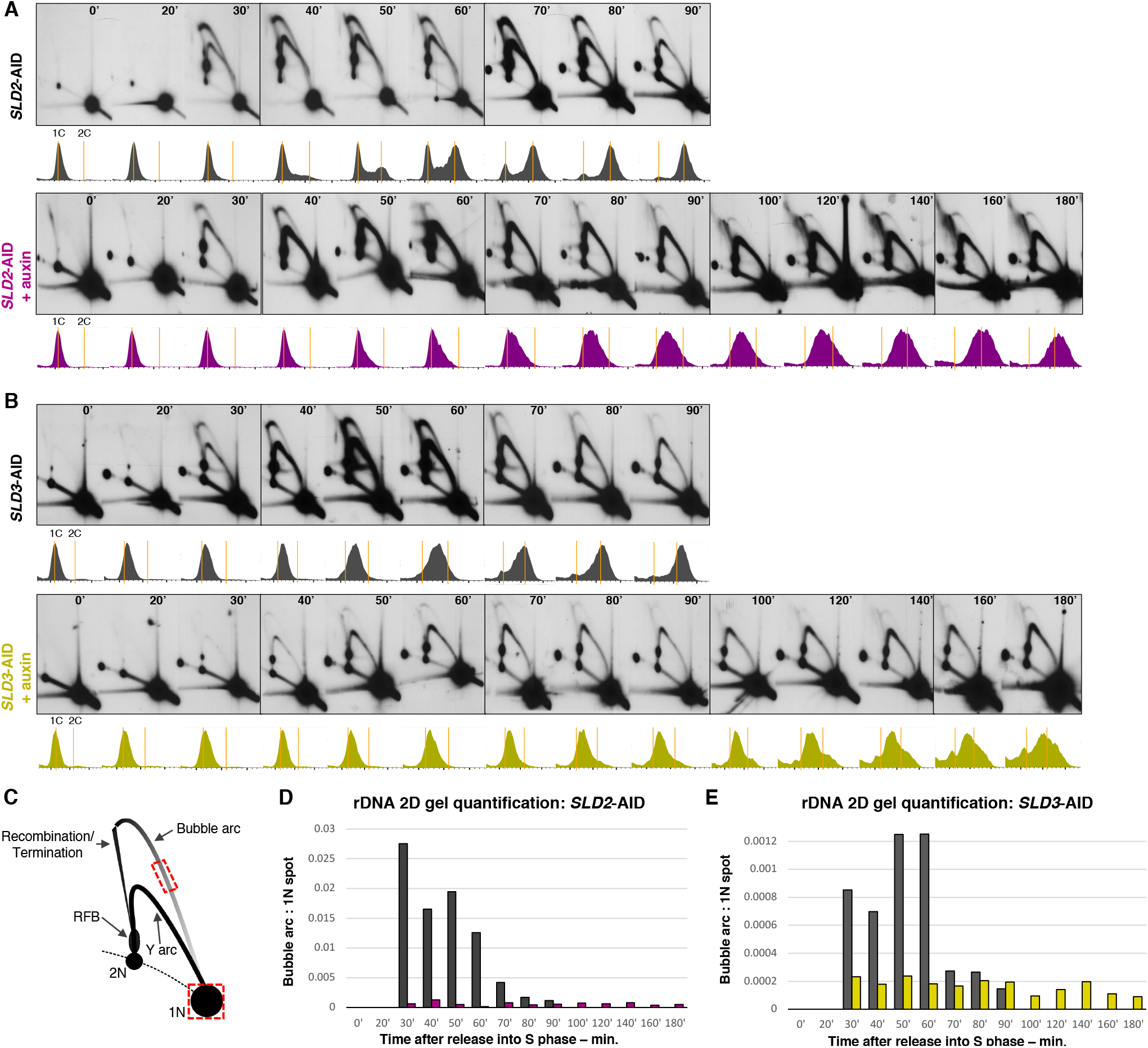
2D gel analysis of the rDNA origin of replication during synchronous S phase. (A) *SLD2*-AID rDNA 2D gels. The resulting Southern blots of Nhel-digested genomic DNA were probed for the non-transcribed spacer region, *NTS2*, which contains the potential rDNA origin. The time relative to release from α factor arrest is indicated in the top corner of each panel. Flow cytometry profiles corresponding to each timed sample are below each panel. Orange lines mark cells with 1C and 2C DNA content. (B) *SLD3*-AID rDNA 2D gels. Sample collection times and FACS profiles are labeled as in (A). (C) Illustration of an rDNA 2D gel and quantification strategy. The red boxes indicate the portions of the bubble arc and the 1N spot used to determine the bubble:1 N ratio. (D) Quantification of the timed *SLD2*-AID rDNA 2D gels. We did not calculate values for T0 and T20 since there was so little bubble arc hybridization signal in those samples. (E) Quantification of *SLD3*-AID rDNA 2D gels as described in (D).

Sld2-depleted cells had reduced bubble:1N ratios throughout S phase with initiation occurring most robustly from T30 to T70 (Figure 3D). Thereafter initiation fell to much lower levels, but bubble signal persisted throughout the lengthened S phase (Figure 3A, D). While termination structures arising from the replication fork barrier (RFB) were present in each S phase sample, from T120 - T180 they became a very prominent species of branched molecule (Figure 3A). In the uninduced control, a modest increase in termination structures occurred at T70 (Figure 3A). The rDNA origin initiation defect in the Sld3-depleted cells was more severe than the defect observed during Sld2 depletion. Bubbles were rare at T30 and remained at a very low level over the entire elongated S phase (Figure 3B, E). There was no discernable increase in termination structures suggesting that initiation events were so widely spaced that by T180, converging forks were not yet close enough to one another to be present on the same Nhe1 fragment.

These rDNA 2D gel data are consistent with a reduction in rDNA origin efficiency without a change in the time at which the origin is first able to fire. The persistent termination structures observed in Sld2-depleted cells are also consistent with the problems in completing rDNA locus replication observed in our CHEF gel analysis. The near-absence of termination structures during Sld3 depletion and the more severe reduction in origin firing efficiency are also consistent with the incomplete rDNA replication deduced from CHEF gel analysis.

### Depletion of Sld2 or Sld3 reduces origin efficiency genome-wide in early S phase

While there is a clear defect in initiation at the rDNA origin when Sld2 or Sld3 is depleted, our CHEF gel analysis also indicates that replication of all chromosomes takes longer. One possible explanation for slowed chromosome replication is that replication forks are slower, although it is difficult to explain how a lack of initiation factors that do not travel with the replication fork would slow the rates of polymerization. To us, it seemed more likely that origins outside of the rDNA are also sensitive to levels of Sld2 and Sld3 and thus have reduced firing efficiencies or delayed times of initiation. In this case, each cell would fire fewer origins and therefore have fewer active replication forks. Consequently, those fewer active forks would need to travel farther to ensure complete chromosome replication and the time required to duplicate the genome would increase. We entertained two models, which are not mutually exclusive, for how unique genomic origin use might be altered during Sld2- or Sld3-depleted S phase: (1) that depletion of Sld2 or Sld3 reduces origin firing efficiency uniformly across the genome, and (2) that depletion of these factors delays the firing of specific origins. In both cases, forks from active origins should be able to travel further after depletion of Sld2 or Sld3 because there are fewer forks active across the genome (Figure 4A) (Zhong et al., 2013).

**Figure 4.**
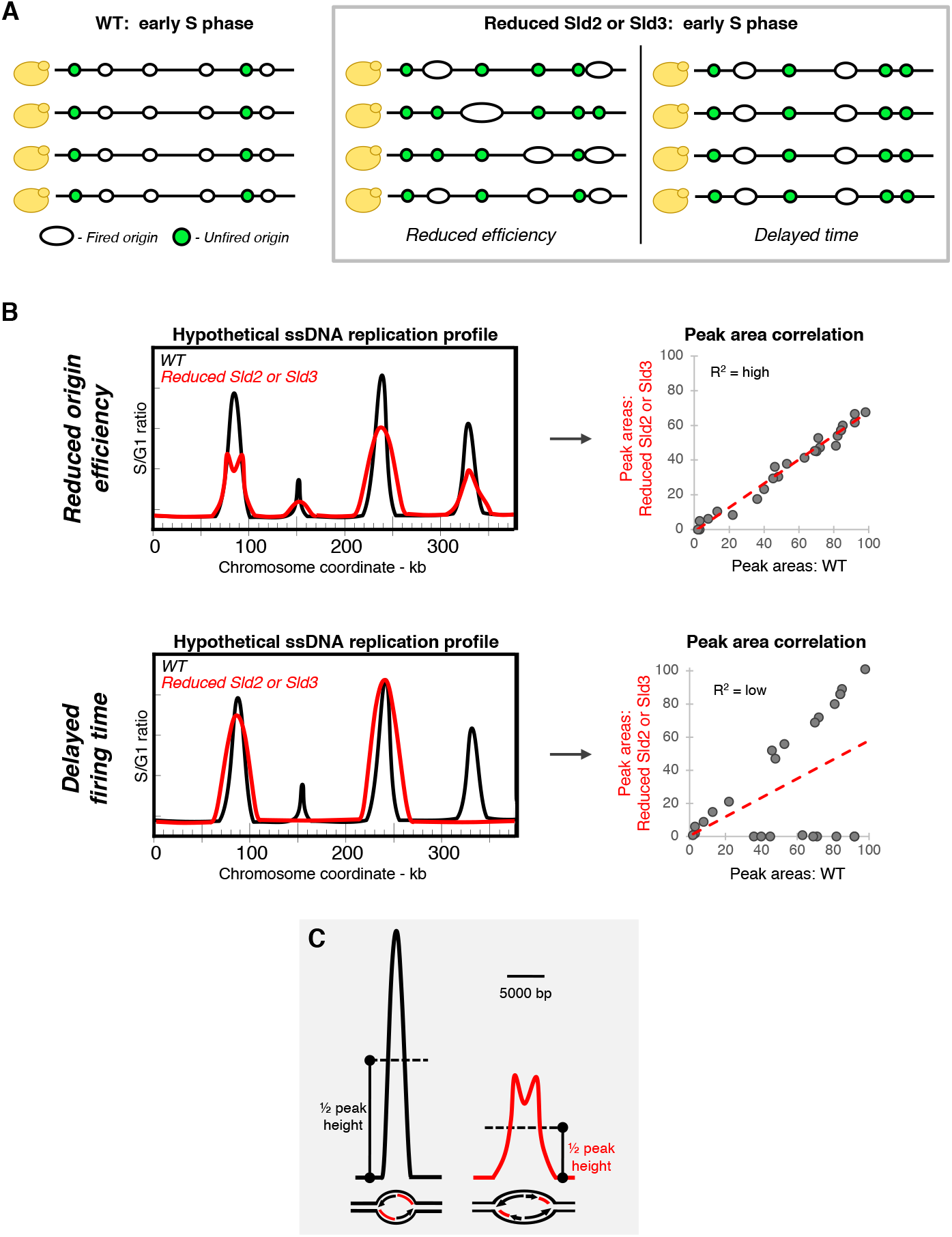
Models for changes to replication initiation during reduced Sld2 or reduced Sld3 S phase. (A) Illustration of predicted changes to early S phase origin firing across a population of cells. If initiation were generally decreased, then compared to a WT population of cells in which four out of six origins fire early in S phase, the same set of origins would be active. Alternatively, specific early origins may be unable to initiate replication until later in S phase, producing two distinct classes of early origins. In both cases, because fewer origins have fired per cell, the forks that are active will move farther before being restricted by the HU treatment (Zhong et al., 2013). (B) Changes to firing time and firing efficiency of early origins measured using a genome-wide assay for ssDNA accumulation at replication forks during S phase in hydroxyurea. Hypothetical ssDNA profiles for WT (black) or Sld2- or Sld3-depleted cultures (red) are shown. If there were a widespread reduction in origin efficiency, we would detect ssDNA peaks at the same locations as in WT. By calculating the area under curve at each origin we can determine the correlation between the sets of peak area values for the two conditions (shown to the right of the hypothetical profiles). A widespread reduction in efficiency would allow for greater replication fork movement genome-wide. If two replication forks are able to move far enough away from each other, the single stranded gaps at the two forks are resolved into separate peaks that flank the position of the origin (the leftmost peak on the hypothetical reduced efficiency profile) (Feng et al., 2011). If depletion of Sld2 or Sld3 resulted in the delay of specific origins until later in S phase, some ssDNA peaks would be absent or extremely reduced in peak height. Since the area under the curve at those delayed origins would be very low compared to WT, the two sets of origin peak area values would be less well-correlated. Individual origins that fail to fire would be identified as data points near 0 on the Y-axis. (C) Confirmation of the reduction in origin efficiency by an increase of the ssDNA peak width at half maximum peak height (Sheu et al., 2014). The black peak is an example of a WT ssDNA origin peak with its corresponding replication bubble and corresponding fluorescent-labelled ssDNA (red) illustrated below the peak. The red peak is at the same origin, but when origin efficiency is reduced. Depending on the synchrony of any particular early origin in the S phase samples, the broader peaks may or may not be “split”.

To examine the activity of non-rDNA origins, we used a single-stranded DNA mapping assay to detect enrichment of ssDNA at replication forks genome-wide by microarray (Feng et al., 2006, 2011). The ssDNA assay allowed us to differentiate between our two models by determining whether or not the same set of origins are active when Sld2 or Sld3 is depleted, as well as to detected changes in replication fork movement that reflect reduced origin efficiency (see Figure 4B and 4C for more details). For this method, we released the cells into S phase in hydroxyurea, which limits dNTP synthesis, and thus limits the timescale of our analysis to early-S phase origins.

We first confirmed that the uninduced degron strains initiated DNA replication from the same origins as WT cells (Figure 5A and Supplemental Figure 4A, B). We then examined the ssDNA profiles for Sld2 and Sld3 depleted conditions. Visual examination of the *SLD2*-AID ssDNA profiles suggested that the same set of early-firing origins are active even when Sld2 has been depleted (Figure 5A), a finding that argues against the possibility that Sld2 or Sld3 depletion delays the firing of specific origins. A comparison of the origin peak areas for the two conditions confirmed this observation (R^2^ = 0.9482) (Figure 5B). Since the same origins are active in the population of cells but S phase progression is impaired, we concluded that origin efficiency must be reduced. The peaks on the *SLD2*-AID plus auxin profiles appeared slightly wider and some were resolved as split peaks (Figure 5A; Supplemental Figure 5). We found a significant difference in replication fork movement by comparing the distributions of origin peak width (Wilcoxon rank sum test p-value < 2.2x10^-16^) (Figure 5C). Overall, these findings are consistent with our first model in which a reduction in origin efficiency across the genome results in an increase in the distance that forks are able to travel when dNTP supply is limited by HU.

**Figure 5.**
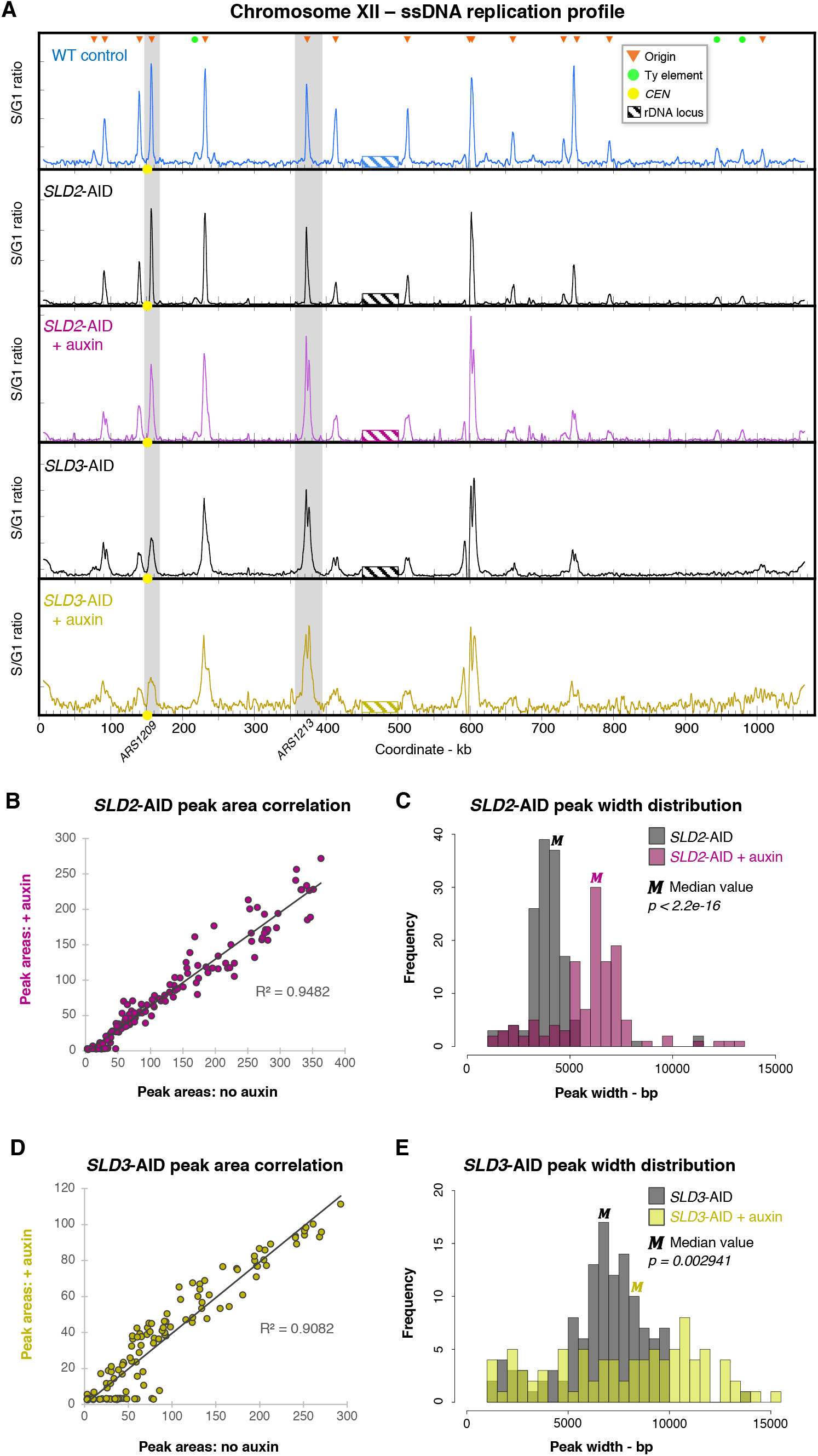
Early S phase replication profiles when Sld2 or Sld3 are depleted. (A) Chr XII replication profiles for WT T30 and T60 for *SLD2*-AID and *SLD3*-AID. Profiles are generated by measuring the ratio of ssDNA labeled in S phase relative to ssDNA labeled in G1. We grew the WT cells in YC + 2% glucose and observed a faster transition to S phase compared with the degron strains that were grown in raffinose. To adjust for this difference, we compared the 30-minute WT profiles to the 60-minute profiles from the AID strains. Orange triangles mark positions of WT origins with a maximum S/G1 ssDNA ratio that is at least 3 SD above average. Green dots with elevated S/G1 ssDNA correspond to regions adjacent to Ty elements that are not origins of replication. Yellow dots on the X axis mark the centromere. Values at the rDNA locus (striped box at coordinates 450-500 kb) are excluded because of low probe density on the microarrays. Two origins, *ARS1209* and *ARS1213*, that we subsequently analyzed by 2D gel (Figure 6B) are highlighted. (B) ssDNA peak area correlation for the two *SLD2*-AID conditions. (C) Distribution of origin peak width for *SLD2*-AID. Widths are calculated in 500 bp intervals. The median peak width was 4000 bp for uninduced *SLD2*-AID and 6000 bp for auxin-treated *SLD2*-AID. Some peak widths were rounded down to zero and therefore excluded from the distribution analysis. An unpaired Wilcoxon Rank Sum test demonstrated a significant difference between the two distributions (p < 2.2x10^-16^). (D) Origin peak area correlation for *SLD3*-AID. (E) Peak width distribution for *SLD3*-AID. The uninduced *SLD3*-AID median was 7000 bp and auxin-treated *SLD3*-AID median was 8500 bp. We excluded zero values before using an unpaired Wilcoxon Rank Sum test to test for significance (p = 0.002941).

**Figure 6.**
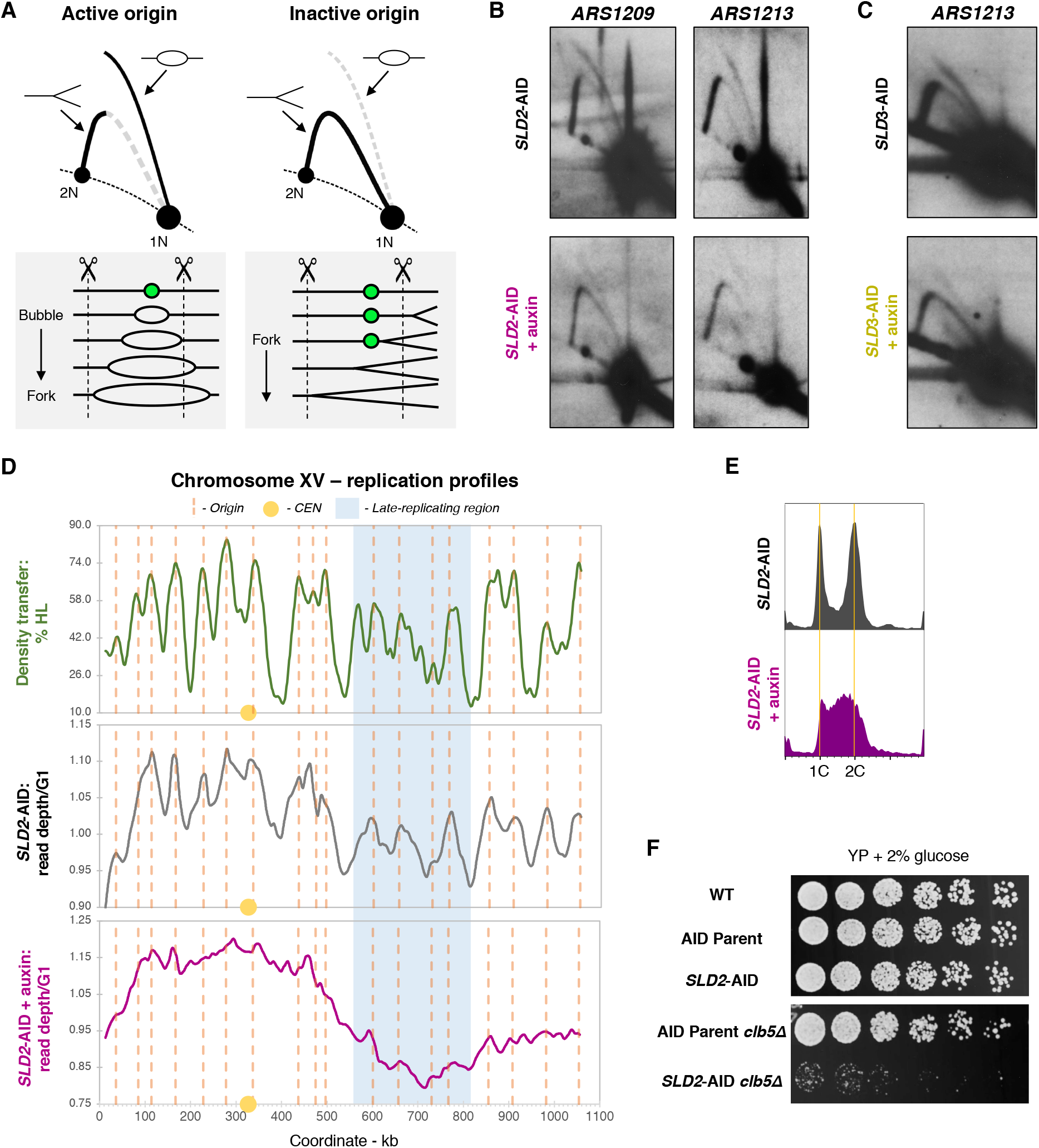
Confirmation of reduced origin efficiency by 2D gel electrophoresis and analysis of late origin activity. (A) Illustration of expected 2D gel signal at an active vs. inactive origin. The positions of bubble and Y intermediates are marked by arrows. The line diagrams below the 2D gel diagrams illustrate how replication forks move through the hypothetical restriction fragment (delimited by the dotted lines) containing an asymmetrically-located potential origin (closed green circle). At active origins, the bubble arc signal predominates and converts to large Ys as one of the forks passes the closest restriction site. However, when the origin fails to fire, the restriction fragment generates a uniform Y arc as forks from an adjacent origin pass through the fragment. (B) *ARS1209* and *ARS1213* 2D gel analysis for *SLD2*-AID. (C) *ARS1213* 2D gel analysis for *SLD3*-AID. (D) Whole genome sequencing (WGS)-based replication profiles for Chr XV in *SLD2*-AID. A replication profile for Chr XV based on WGS read-depth for untreated *SLD2*-AID is shown in the center panel and compared to our previously published replication profile determined from a density transfer-to-microarray experiment (top panel; replotted from Alvino et al., 2007). The vertical dotted orange lines correspond to confirmed origins in OriDB (Siow et al., 2012) that are detected in both the density transfer and the uninduced *SLD2*-AID WGS profiles. The yellow circle marks the centromere and the region highlighted in blue indicates the late-replicating domain. The bottom panel shows the WGS read-depth profile for the *SLD2*-AID strain treated with auxin for two hours. (E) Flow cytometry profiles for asynchronous *SLD2*-AID collected for whole genome sequencing. Orange lines indicate 1C and 2C DNA cells. The two-hour auxin treatment of *SLD2*-AID cells causes more than 50% of the cells to be distributed throughout S phase. (F) Spot test of *SLD2*-AID *clb5Δ* strain growth and viability. Cell cultures were serially diluted 1:3, spotted onto YEPD, grown at 30°C for three days and then photographed.

We applied the same analysis to the *SLD3*-AID strain and also observed peaks at the same origins in the two conditions (Figure 5A; Supplemental Figure 5). The subsequent genome-wide comparison of peak area values revealed high correlation (R^2^ = 0.9082) confirming that the same origins are active even when Sld3 is depleted (Figure 5D). We observed that the origin peaks in uninduced *SLD3*-AID cells were much broader and were often split compared to WT. This unusual peak topology suggested that the uninduced *SLD3*-AID strain had reduced origin efficiency, a finding that is in agreement with our previous observations of a delayed start of S phase, slowed Chr XII replication, and reduced rDNA copy number in the uninduced strain. Adding auxin to further reduce Sld3 function did correspond with an additional increase in replication fork movement (Wilcoxon rank sum p-value = 0.002941) (Figure 5E). We attribute the less drastic change in fork movement upon Sld3 depletion compared to Sld2 to partial loss of Sld3 function caused by the degron tag. To test for compromised Sld3 function, we constructed a version of the *SLD3*-AID strain without the E3 ubiquitin ligase. We found that both isolates of this strain had a larger population of S phase cells than WT (Supplemental Figure 6A) and that the presence of the AID tag correlated with reduced rDNA origin efficiency (Supplemental Figure 6B). Together, these data demonstrate that the degron tag on Sld3 reduces its function and, therefore, origin efficiency.

### 2D gel electrophoresis of individual origins confirms reduced efficiency

We concluded that the ssDNA mapping assay data primarily support the model in which Sld2 or Sld3 depletion reduces origin efficiency genome-wide. To confirm reduced efficiency at non-rDNA origins, we analyzed the unique early origins *ARS1209* and *ARS1213* by 2D gel electrophoresis (Figure 6A). We isolated genomic DNA from asynchronously growing cells that we treated with auxin for two hours. We observed striking changes in the cell cycle profiles indicating that cells had accumulated in S phase: more than 50% of auxin-treated *SLD2*-AID cells were distributed across S phase and the majority of auxin-treated *SLD3*-AID cells had accumulated in early S phase (Supplemental Figure 7A). We found a reduction in replication bubble signal and an increased Y signal intensity for both origins in the Sld2-AID strain exposed to auxin (Figure 6B). *ARS1213* efficiency was also lower following Sld3 depletion (Figure 6C), although the *ARS1209* gel was less conclusive (Supplemental Figure 7B). Finally, we tested efficiency at late-firing *ARS522* and found that depletion of either Sld2 or Sld3 diminished its efficiency (Supplemental Figure 7C).

### Late-firing origins are still capable of firing when origin efficiency is reduced genome-wide

One of the predictions of the Mantiero et al. model (2011) is that the temporal program of origin firing is carried out by recycling the limiting factors to later-firing origins. We considered the possibility that if recycling of Sld2 or Sld3 were prevented by auxin-induced degradation, late origins may be rendered inactive. Based on our 2D gel analysis of late-firing *ARS522* in which initiation signal was below the level of detection, it seemed possible that reduced efficiency resulting from initiation factor depletion suppressed initiation more for late-firing origins than for early-firing origins. To query late origin activation across the genome we used whole genome sequencing (WGS) of asynchronous cells to profile DNA replication (Müller et al., 2014). In an asynchronous culture it is possible to detect origin activation as spikes in copy number relative to adjacent non-origin sequences (Supplemental Figure 8A). Since *SLD3*-AID cells failed to proceed past early S phase after 2 hours in auxin, we limited our analysis to the *SLD2*-AID strain which is enriched in cells across S phase (Supplemental Figure 7A) and compared the resulting replication profiles with the non-induced culture. This method allowed us to survey origin firing for all of S phase (Supplemental Figure 9A).

In the absence of auxin, we found clear peaks in copy number amplitude closely paralleling the results from our previous study using density transfer and microarray analysis of synchronized wild-type cells (Figure 6D). Chromosome XV illustrates the distinction between early- and late-firing origins since it contains an internal late-replicating region that we previously showed is due to a cluster of late-firing origins (blue box in Figure 6D) (McCune et al., 2008; Raghuraman et al., 2001). In the presence of auxin, *SLD2*-AID maintains both the early- and late-replicating domains but has lost much of the clear definition between origin peaks (Figure 6D). However, the distinctions between origins in both early- and late-replicating regions are diminished uniformly, a feature that is found genome wide (Supplemental Figure 9B). Because we still observe small spikes in read depth at nearly all origins when Sld2 is depleted, regardless of firing time, we conclude that late origins are active and do not experience a greater reduction in firing efficiency than the early origins.

To confirm that late origin activation still occurs when Sld2 is depleted, we deleted *CLB5*, which is needed for origin firing in late S phase (Donaldson et al., 1998). If the effects of *CLB5* deletion on growth rate, cell cycle progression, or viability worsen the effects of Sld2 depletion alone, we could conclude that origins are still able to fire in late S phase when Sld2 is depleted. Deletion of *CLB5* in the AID parent strain had a negligible impact on growth (Figure 6F). However, our attempts to delete *CLB5* in *SLD2*-AID, both by gene replacement using standard lithium acetate transformation and by crossing *SLD2*-AID into the AID parent *clb5Δ* strain, were of limited success. We isolated and confirmed a single, sick *SLD2*-AID *clb5Δ* clone. The single *SLD2*-AID *clb5Δ* isolate could not be grown to high enough culture density for any experiments besides a spot-test, which confirmed its growth defect even without the addition of auxin (Figure 6H). The *SLD2*-AID *clb5Δ* clone was inviable when exposed to 200 mM HU (Supplemental Figure 9). These results indicate that when Sld2 activity is compromised, the cells rely on *CLB5* and therefore late origin firing to complete S phase.

## Discussion

Inspired by the observation that increasing the cellular abundance of four limiting initiation factors advances origin firing time and enhances origin efficiency (Mantiero et al., 2011), we wanted to ask: Do these factors contribute differentially to origin efficiency and timing? Would further restricting the availability of these factors have disproportionate consequences for specific origins and cause instability at particular regions of the genome?

We find that the abundance of the limiting DNA replication initiation factors Sld2 and Sld3, individually, contribute primarily to origin efficiency, and that the effects are global rather than restricted to particular genomic regions or classes of origins. When we induce depletion of either factor, both the rDNA and unique origins experience a reduction in efficiency, while the onset of initiation does not change. The rDNA 2D gel analysis, in which we were able to measure the time of initiation and origin efficiency in a single experiment, demonstrates clearly that reduced origin efficiency does not necessarily correspond with a delay in the onset of origin firing. Using the ssDNA replication assay, we infer from the genome-wide increases in replication fork movement that the same relationship between origin efficiency and time of initiation holds true outside of the rDNA array. The high correlation between origin peak areas for uninduced and induced *SLD2*-AID and *SLD3*-AID cells reiterates that origin selection does not change in response to initiation factor depletion and that no subset of origins is more affected than others.

The uniform impact on origin firing across the genome is also demonstrated by our marker-frequency analysis from WGS; we find that both early and late origins suffer equivalently from depletion of Sld2. These data are supported by the synthetic sickness of *SLD2*-AID with the absence of the Clb5 cyclin, the CDK subunit that is required for late origin activation. To our knowledge, the only function common to Sld2 and Clb5 is origin activation; we therefore conclude that the synthetic sickness stems from difficulty in completing genome replication. We interpret these results as evidence that even when origin efficiency is extremely low, origin activation still occurs in late S phase, as these cells evidently rely upon origins firing later in S phase for genome replication. We suspect that the combination of the degron tag on Sld2 and the restricted window for S-CDK activity in a *clb5Δ* background lowers the levels of functional Sld2 during late S phase.

The consequence of lowered efficiency of origin firing across all of S phase is a blurring of the distinction between origins and their intervening termination regions. In the WGS replication profiles from uninduced *SLD2*-AID, origins and termination regions are distinguished as high peaks and low valleys, respectively. When Sld2 is depleted, there is a loss of distinction between origin peaks and the broadest features of replication remain—higher read depth in the early-replicating regions and lower read depth in the late-replicating regions. This WGS profile is strikingly similar to the megabasepair-sized “timing domains” observed in replicating metazoan genomes (Cayrou et al., 2011; Petryk et al., 2016; Rhind and Gilbert, 2013). In metazoans, it is thought that these timing domains contain many adjacent, low efficiency origins that tend to fire within the same span of time during S phase (Petryk et al., 2016; Rhind and Gilbert, 2013).

The widespread reduction of origin efficiency in uninduced *SLD3*-AID, due to partial loss of function of the tagged protein, also supports our conclusions regarding the control of origin efficiency. Even without inducing depletion of Sld3, the degron domain at the C-terminus of Sld3 impairs origin firing both at the rDNA and unique chromosomal origins. We suspect that the reduced rDNA copy number in *SLD3*-AID is a compensatory change that mitigates the susceptibility of rDNA stability to low origin efficiency—a specific source of instability that we observed by CHEF gel analysis during Sld2- or Sld3-depleted S phase. While origin efficiency is compromised in uninduced *SLD3*-AID, the same set of early origins fires, but at reduced levels across the population of cells.

It occurs to us that our observations about the relationship between origin efficiency and firing time could inform an ongoing debate in the field regarding a model for replication initiation known as the “stochastic model of origin firing.” According to this model, origin firing time is simply a consequence of origin efficiency and the two features are determined by the same underlying molecular mechanisms (de Moura et al., 2010; Rhind, 2006; Yang et al., 2010). A key aspect of this model is that any features that increase the probability that an origin will fire will likewise advance its time of replication (Rhind, 2006). As a corollary, this model predicts that any treatment that decreases an origin’s efficiency would likewise cause a delay in its time of initiation. In contrast to our conclusions that Sld2 and Sld3 impact origin efficiency independent of origin firing time, this model would predict that any changes to SSDD factor abundance would impact both origin efficiency and time of origin firing.

In fact, our study reinforces the notion that origin firing efficiency and time of origin firing are separable. We propose that most experimental evidence, including ours, points to a more complex relationship among features of the genome that can influence time and efficiency through separate mechanisms. For example, variable replication time across a genome is observed even when origin efficiency is increased: during overexpression of the SSDD factors, Mantiero et al., (2011) found subtle differences in replication time at multiple loci in the yeast genome. A recent timing study of pre-MBT zebrafish embryos demonstrated that, even during the rapid cell divisions when origin efficiency is high across the entire genome, there are still early- and late-replicating regions of the genome (Siefert et al., 2017). Our observed changes in replication efficiency independent of origin firing time in response to Sld2 or Sld3 depletion are in accordance with a model for variation in replication activity in which origin efficiency and time of firing are influenced by many different features of the genome.

While Chr XII-specific destabilization under reduced-initiation conditions has been observed previously (Ide et al., 2007; Salim et al., 2017; Sanchez et al., 2017), our observations of rDNA array instability suggest a slightly different interpretation of a similar phenomenon described by Ide et al. (Ide et al., 2007). After shifting asynchronously-growing *orc1-4* cells to the non-permissive temperature, Ide et al. (2007) reported that instability of Chr XII preceded instability of the other chromosomes. Primarily from this difference in time, they proposed that the rDNA array acts as a sensor for problems with DNA replication initiation and that instability of the rDNA acts as something of a “canary in a coal mine” for the rest of the genome. No mechanism for transduction of such a signal was identified in their work, and the point in the cell cycle at which they first observed instability is unclear—in their study, Ide et al. (Ide et al., 2007) did not synchronize cells in S phase, so the time of onset of Chr XII instability was not determined. If the rDNA is the first part of the genome to experience initiation defects and, based on sensing those problems, it then relays that information to the rest of the genome, one would predict that replication initiation defects would be detected on Chr XII before the other chromosomes. However, in our synchronous S phase CHEF gel analysis, incomplete replication of Chr XII is not evident any earlier than the effects of reduced initiation detected on other chromosomes (compare T0 - T60 quantification for all chromosomes in Supplemental Figure 2). We conclude that Chr XII is inherently less stable than other chromosomes due to the array of identical origins that spans more than 2 Mb of DNA. Furthermore, the instability of Chr XII has no bearing on the stability of the other chromosomes; in our CHEF gel experiments, we did not detect instability of other chromosomes during the synchronous S phase or the after the cells had lost synchrony.

The rDNA instability in *S. cerevisiae* suggests that large spans of repeated DNA elements may likewise be a problem for genomes with much more repetitive content, such as the human genome. For example, the rDNA loci in humans, while divided into arrays on 6 different chromosomes, also have a tandem repeat structure (Stults et al., 2008). Human centromeres, which are megabasepairs long, also consist of tandem repeats primarily made up of α satellite DNA (Melters et al., 2013; Willard, 1990). The yeast rDNA locus is the only part of the yeast genome that approximates the structure of these long, tandemly-repeated parts of the human genome, and its sensitivity to replication initiation defects suggests that defective replication initiation may also impact the stability of similarly repetitive regions of metazoan genomes. Generally, it would be challenging to determine whether large repeated DNAs in metazoan cells are similarly sensitive to replication initiation defects because the requirements for origins are still unclear in those organisms (reviewed comprehensively by Prioleau and MacAlpine, 2016). Nevertheless, our work suggests that DNA replication through repetitive sequence could be a potential challenge for metazoans in terms of ensuring complete replication and maintaining genome stability.

## Materials and Methods

### Strain construction

The parental strain for this study is W303 *rad5Δ MATa*, which is used as a wild-type (WT) control throughout. We used standard yeast lithium acetate transformation for all transformations. In the auxin inducible degron (AID) strains, the *GAL1-10-*Os*TIR1*-9myc construct is integrated at *URA3*. We C-terminally tagged Sld2 and Sld3 with 3xmini-AID by transformation and confirmed integration of the AID tag by PCR and Southern blot hybridization. Takashi Kubota and Anne Donaldson at the University of Aberdeen generously donated the auxin degron construct plasmids to us. We confirmed Sld2 and Sld3 degradation following auxin induction by flow cytometry for delayed or slowed S phase. We constructed the *RAD53*-2xHA tagged versions of the AID strains by integrating linearized plasmid containing a partial copy of *RAD53* with the HA tag. The Bedalov lab at the Fred Hutchinson Cancer Research Center donated the *RAD53*-2xHA plasmids. See Supplemental Table 1 for oligos and Supplemental Table 2 for list of strains.

### AID strain induction

All liquid culture was in yeast synthetic (YC) medium. For asynchronous experiments we grew cells in YC + 1% raffinose (YCR) medium until early log phase and induced *Os*TIR1-9myc expression with 2% galactose for 3 hours before adding 0.5 mM auxin (IAA) for a minimum of 2 hours. Unless specified, negative controls are the degron strain cells without auxin (see Figure 1B). As needed, we diluted cultures during galactose induction to maintain log phase growth. For synchronous S phase experiments, we grew cultures in YCR to early log phase, then synchronized them with 3 μM α-factor for 3.5 hours. Thirty minutes after adding α-factor, we added 2% galactose and, one hour before release, we split the cultures and added auxin to one half at a final concentration of 0.5 mM. We collected flow cytometry samples for every experiment to confirm initiation factor degradation.

### Flow cytometry

We collected one mL of culture per sample and prepared samples as in Alvino, et al (2007) using SYTOX Green Dead Cell Stain (ThermoFisher). We assayed 10,000 events per sample on a BD FACSCanto cytometer and analyzed the data in FlowJo. For the degron strains, we gated all samples based on the uninduced asynchronous culture.

### Growth and viability

To determine viability, we plated 300 cells in triplicate on YEPD and counted colonies after 2 days of growth. For spot-testing, we suspended the cells in water, sonicated, then serially diluted the cells in a 1:3 ratio and spotted 5 μl onto appropriate plates. We grew the cells at 30°C and photographed the plates each day.

### Western blotting

For each sample, we collected ~4.0 x 10^7^ cells at mid-log phase. The positive control samples were treated with 0.1% MMS for 1 hour. We extracted protein by bead beating in SUMEB pH 6.8 (1% SDS, 8 M Urea, 10 mM MOPS pH 6.8, 10 mM EDTA, and 0.01% bromophenol blue) plus ThermoFisher protease inhibitor cocktail and 5% 2-Mercaptoethanol. We probed with 1:2500 α-HA (Abcam #ab128131) and 1:5000 α-GAPDH (Abcam #ab9385) as a loading control.

### CHEF gel analysis of chromosome stability

We used contour-clamped homogeneous electric field (CHEF) gel electrophoresis to assay chromosome integrity and stability during synchronous S phase (Cha and Kleckner, 2002; Feng et al., 2011; Hennessy et al., 1991). We collected 5 mL of culture for each timed sample then embedded the cells in agarose. We ran the samples on a 1% agarose gel on Bio-Rad CHEF electrophoresis apparatus at 100 V for 68 hours, switch time ramped from 300 to 900 seconds as in Kwan et al. (2016). We transferred the gels using standard Southern blotting methods to GeneScreen nylon membrane (PerkinElmer) and probed the same blots multiple times for single copy sequences on different chromosomes, stripping between each hybridization. We used a Bio-Rad Personal Molecular Imager and QuantityOne software to quantify hybridization signal.

For the *in gelo* I-PpoI restriction digest, we washed the plugs 3x in 10 mM Tris, then pipetted 75 μl I-Ppo1 buffer plus 0.5 μl I-Ppo1 onto a small slice of each plug and incubated for 1 hour at 37°C. We ran each sample using the same CHEF gel parameters as above.

### 2D gel electrophoresis

We performed 2D gel electrophoresis analysis for the high copy rDNA origin and unique origins using different DNA purification and restriction digest protocols for each type. For each timed rDNA sample, we collected 50 ml culture volume by chilling the cells on frozen 0.1% sodium azide plus 0.025 M EDTA. We then embedded the cells in 0.5% low melt (SeaPlaque) agarose plugs and spheroplasted (Feng et al., 2011). To digest the DNA, we cut a single plug in half and washed it 3x in 10 mM Tris, then equilibrated 3x in 200 μl NEBuffer 2.1 on ice with gentle shaking. We removed the buffer and pipetted 3 μl NheI (10,000 units/ml; NEB) directly onto the plug and incubated it for 4 hours at 37°C. We loaded each plug onto a comb and poured the 0.5% agarose first dimension gel around the plugs. We ran the first-dimension gel for 20 hours at 1V/cm without ethidium bromide. We cut the resulting gel slice at least 1 cm below the 4.7 kb NheI fragment that contained the origin, then transferred each gel slice to a casting tray and poured the 2^nd^ dimension gel. We ran the 1.1% agarose second-dimension gel with 0.3 μg/ml ethidium bromide for 6 hours at 7 V/cm in the cold room with circulating buffer. We probed the membranes with the *NTS2* sequence and quantified the rDNA 2D gels as depicted in Figure 3C. For single copy origins, we collected 300 mL of mid-log phase culture (OD_660_ 0.5 – 0.7). We chilled the sample on frozen sodium azide and EDTA and purified genomic DNA using a variation of the “NIB (nuclear isolation buffer) & Grab” DNA purification protocol detailed on the Brewer/Raghuraman lab website (Payen et al., 2013). Instead of using glass beads and vortexing alone to lyse the cells, we incubated the cells in 1 mg/ml zymolyase in NIB for 30 minutes at room temperature, and then used one 30s round of glass bead-beating to ensure lysis before continuing with the standard NIB & Grab protocol. We find that zymolyase-mediated cell lysis better preserves replication intermediates.

We used all of the DNA purified from 300 mL of culture (a few micrograms) for each 2D gel of one origin. We used the enzyme BanI to analyze *ARS1209* (4.8 kb fragment) and *ARS1213* (3.7 kb fragment). Due to a polymorphism that changed the BanI site in *SLD3*-AID, we used BspHI to analyze *ARS1209* (4.9 kb fragment). For *ARS522*, we used XbaI to cut a 3.2 kb origin fragment. We ran the 0.4% first dimension agarose gel at 1 V/cm for 18 hours without ethidium bromide. We ran the 1.1% agarose second-dimension gel with 0.3 μg/ml ethidium bromide in the cold room with buffer circulation at 7 V/cm for 4 – 5 hours, depending on the size of the origin fragment. After transfer, we probed the blots with a fragment containing the origin of interest. See Supplemental Table 1 for the oligos used to amplify the probe fragments.

### ssDNA mapping replication assay

We grew cultures to OD_660_ = 0.25, α-factor arrested them and added auxin as described above. We grew the WT cells in YC + 2% glucose. We collected S phase samples at 30, 60, 90, and 120 minutes after release. We prepared samples using the *in gelo* ssDNA labeling protocol detailed by Feng et al. (2011), and used β-agarase (NEB #M0392S) to recover the labeled ssDNA. We co-hybridized one G1 and one S phase sample to Agilent ChIP-on-chip 4x44 *S. cerevisiae* microarray. We processed and normalized the array data as described in Feng, et al. (2006), and used LOESS regression to smooth the data over a 6 kb window. Raw intensity values and processed microarray data are available in Supplemental Data Files 1-5.

### Origin peak area and peak width at half maximum height

We used a custom Python script to calculate origin peak areas and peak width at half the maximum peak height for each of the origins detected in the ssDNA assay data (Sanchez et al., 2017). We used the list of known origins from the DNA Replication Origin Database OriDB (http://cerevisiae.oridb.org/) (Siow et al., 2012). Since not all origins fire during S phase in HU, using the complete OriDB list resulted in “overcalled” peaks, so we reduced the origins list based on the WT control. We included origin peaks with a maximum height more than 3 standard deviations above the average S/G1 ssDNA ratio in WT. Additionally, we noticed that at some peaks, more than a single origin is identified in the database, so we also excluded multiple calls at a single origin. To confirm these cutoffs, we compared the peak areas from uninduced *SLD2*-AID with WT, as well as uninduced *SLD3*-AID with WT (Supplemental Figure 4). Based on the high correlation between WT and the uninduced degron strains, we used this pared-down origin list for all subsequent analysis. Supplemental Table 3 includes the full list of origins analyzed by ssDNA mapping. We found that even though the microarray probes for Ty elements are excluded during data processing, there can be artifactual labeling of ssDNA at Ty-adjacent loci that generates ssDNA peaks. Since those peaks do not reflect origin activity, we have indicated those loci on replication profiles and listed them in Supplemental Table 4. Finally, we compared the distributions of peak width with an unpaired Wilcoxon rank-sum test in R. In the Python script, peak widths are rounded to the closest 500 bp interval, so some width values are calculated as 0 bp and were thus excluded from the distribution analysis.

### WGS replication profiling

For each sample, we collected 50 mL mid-log phase (OD_660_ ≈ 0.5) cells as well as a G1-arrested control sample after 120 minutes in α-factor. We purified genomic DNA by the standard Smash & Grab protocol (Rose, 1990). We sheared the DNA to an average length of 260 bp using a Covaris ultrasonicator, then treated the sample with 0.01 mg/ml RNAse A before a final clean up using the Zymo Research DNA Clean and Concentrator-25 kit. We end-repaired and adapter-ligated 100 ng of DNA per sample with the KAPA Biosystems Hyper Prep Kit and the KAPA Single-Indexed Adapter Kit. We sequenced on the Illumina NextSeq 500 platform. We aligned single reads of at least 25 bp to Saccer1 using Bowtie2 then binned the reads into 1 kb windows using custom Python scripts. When possible, we aimed for 10 million reads per sample before generating replication profiles.

To generate profiles, we first calculated average read depth for each sample, excluding the rDNA locus, 2-micron plasmid, and the mitochondrial DNA and normalized the read depth in each 1 kb bin to this average. We determined marker frequency for each bin by dividing the bin read depth by the corresponding bin read depth of the G1 cells. We found that poor mappability of repetitive subtelomeric DNA sequences, transposable elements, and highly conserved paralogous genes descended from the whole genome duplication event resulted in outlying marker frequency values that distorted the replication profiles when smoothing to the data. Consequently, we excluded these poorly-mapped regions, which are listed in Supplemental Tables 5 and 6, from each dataset. Finally, we LOESS smoothed the marker frequency values in 50 kb windows to generate replication profiles. All sequencing reads are available at SRA accession #SRP156227. Processed data are included in Supplemental Data Files 6 and 7.

## Acknowledgements

We thank Takashi Kubota and Anne Donaldson for AID strain plasmid constructs and the Bedalov Lab at FHCRC for the Rad53-2xHA plasmid. Thank you to Kate Sitko for western blot advice and reagents as well as to Jason Stephany for assistance with Illumina sequencing. This work was supported by the University of Washington Genome Training Grant T32HG000035, NIGMS R01 GM018926, and NIGMS R35 GM122497.

## References

Alvino, G.M., Collingwood, D., Murphy, J.M., Delrow, J., Brewer, B.J., and Raghuraman, M.K. (2007). Replication in Hydroxyurea: It’s a Matter of Time. Mol. Cell. Biol. 27, 6396–6406.

Bell, S.P., and Labib, K. (2016). Chromosome Duplication in Saccharomyces cerevisiae. Genetics 203, 1027–1067.

Bell, S.P., and Stillman, B. (1992). ATP-dependent recognition of eukaryotic origins of DNA replication by a multiprotein complex. Nature 357, 128–134.

Bloom, J., and Cross, F.R. (2007). Multiple levels of cyclin specificity in cell-cycle control. Nature Reviews Molecular Cell Biology 8, 149–160.

Bousset, K., and Diffley, J.F.X. (1998). The Cdc7 protein kinase is required for origin firing during S phase. Genes & Development 12, 480–490.

Brewer, B.J., and Fangman, W.L. (1988). A replication fork barrier at the 3' end of yeast ribosomal RNA genes. Cell 55, 637–643.

Cairns, J. (1966). Autoradiography of HeLa cell DNA. Journal of Molecular Biology 15, 372–373.

Cayrou, C., Coulombe, P., Vigneron, A., Stanojcic, S., Ganier, O., Peiffer, I., Rivals, E., Puy, A., Laurent-Chabalier, S., Desprat, R., et al. (2011). Genome-scale analysis of metazoan replication origins reveals their organization in specific but flexible sites defined by conserved features. Genome Res. 21, 1438–1449.

Cha, R.S., and Kleckner, N. (2002). ATR Homolog Mec1 Promotes Fork Progression, Thus Averting Breaks in Replication Slow Zones. Science 297, 602–606.

Collart, C., Allen, G.E., Bradshaw, C.R., Smith, J.C., and Zegerman, P. (2013). Titration of Four Replication Factors Is Essential for the Xenopus laevis Midblastula Transition. Science 341, 893–896.

Deegan, T.D., Yeeles, J.T., and Diffley, J.F. (2016). Phosphopeptide binding by Sld3 links Dbf4-dependent kinase to MCM replicative helicase activation. The EMBO Journal 35, 961–973.

Diffley, J.F.X., and Cocker, J.H. (1992). Protein-DNA interactions at a yeast replication origin. Nature 357, 169–172.

Diffley, J.F.X., Cocker, J.H., Dowell, S.J., and Rowley, A. (1994). Two steps in the assembly of complexes at yeast replication origins in vivo. Cell 78, 303–316.

Donaldson, A.D., Raghuraman, M.K., Friedman, K.L., Cross, F.R., Brewer, B.J., and Fangman, W.L. (1998). CLB5-dependent activation of late replication origins in S. cerevisiae. Molecular Cell 2, 173–182.

Douglas, M.E., Ali, F.A., Costa, A., and Diffley, J.F.X. (2018). The mechanism of eukaryotic CMG helicase activation. Nature 555, 265–268.

Ellison, E.L., and Vogt, V.M. (1993). Interaction of the intron-encoded mobility endonuclease I-PpoI with its target site. Mol Cell Biol 13, 7531–7539.

Enserink, J.M., and Kolodner, R.D. (2010). An overview of Cdk1-controlled targets and processes. Cell Div 5, 11.

Farrell, J.A., and O’Farrell, P.H. (2014). From Egg to Gastrula: How the Cell Cycle Is Remodeled During the Drosophila Mid-Blastula Transition. Annual Review of Genetics 48, 269–294.

Feng, W., Collingwood, D., Boeck, M.E., Fox, L.A., Alvino, G.M., Fangman, W.L., Raghuraman, M.K., and Brewer, B.J. (2006). Genomic mapping of single-stranded DNA in hydroxyurea-challenged yeasts identifies origins of replication. Nat Cell Biol 8, 148–155.

Feng, W., Di Rienzi, S.C., Raghuraman, M.K., and Brewer, B.J. (2011). Replication Stress-Induced Chromosome Breakage Is Correlated with Replication Fork Progression and Is Preceded by Single-Stranded DNA Formation. G3 (Bethesda) 1, 327–335.

Ferguson, B.M., Brewer, B.J., Reynolds, A.E., and Fangman, W.L. (1991). A yeast origin of replication is activated late in S phase. Cell 65, 507–515.

Foiani, M., Pellicioli, A., Lopes, M., Lucca, C., Ferrari, M., Liberi, G., Muzi Falconi, M., and Plevani, P. (2000). DNA damage checkpoints and DNA replication controls in Saccharomyces cerevisiae. Mutation Research/Fundamental and Molecular Mechanisms of Mutagenesis 451, 187–196.

Fritze, C.E., Verschueren, K., Strich, R., and Easton Esposito, R. (1997). Direct evidence for SIR2 modulation of chromatin structure in yeast rDNA. EMBO J 16, 6495–6509.

Ghaemmaghami, S., Huh, W.-K., Bower, K., Howson, R.W., Belle, A., Dephoure, N., O’Shea, E.K., and Weissman, J.S. (2003). Global analysis of protein expression in yeast. Nature 425, 737–741.

Hennessy, K.M., Lee, A., Chen, E., and Botstein, D. (1991). A group of interacting yeast DNA replicatlon genes. Genes & Development 5, 13.

Ide, S., Watanabe, K., Watanabe, H., Shirahige, K., Kobayashi, T., and Maki, H. (2007). Abnormality in Initiation Program of DNA Replication Is Monitored by the Highly Repetitive rRNA Gene Array on Chromosome XII in Budding Yeast. Molecular and Cellular Biology 27, 568–578.

Ilves, I., Petojevic, T., Pesavento, J.J., and Botchan, M.R. (2010). Activation of the MCM2-7 Helicase by Association with Cdc45 and GINS Proteins. Molecular Cell 37, 247–258.

Jackson, A.L., Pahl, P.M., Harrison, K., Rosamond, J., and Sclafani, R.A. (1993). Cell cycle regulation of the yeast Cdc7 protein kinase by association with the Dbf4 protein. Molecular and Cellular Biology 13, 2899–2908.

Kamimura, Y., Tak, Y.-S., Sugino, A., and Araki, H. (2001). Sld3, which interacts with Cdc45 (Sld4), functions for chromosomal DNA replication in Saccharomyces cerevisiae. The EMBO Journal 20, 2097–2107.

Koren, A., Handsaker, R.E., Kamitaki, N., Karlić, R., Ghosh, S., Polak, P., Eggan, K., and McCarroll, S.A. (2014). Genetic Variation in Human DNA Replication Timing. Cell 159, 1015–1026.

Kouprina, N.Y., and Larionov, V.L. (1983). The study of a rDNA replicator in Saccharomyces. Current Genetics 7, 433–438.

Kubota, T., Nishimura, K., Kanemaki, M., and Donaldson, A. (2013). The Elg1 replication factor C-like complex functions in PCNA unloading during DNA replication. Molecular Cell 50, 273–280.

Kwan, E.X., Foss, E.J., Tsuchiyama, S., Alvino, G.M., Kruglyak, L., Kaeberlein, M., Raghuraman, M.K., Brewer, B.J., Kennedy, B.K., and Bedalov, A. (2013). A Natural Polymorphism in rDNA Replication Origins Links Origin Activation with Calorie Restriction and Lifespan. PLoS Genetics 9, e1003329.

Kwan, E.X., Wang, X.S., Amemiya, H.M., Brewer, B.J., and Raghuraman, M.K. (2016). rDNA Copy Number Variants Are Frequent Passenger Mutations in *Saccharomyces cerevisiae* Deletion Collections and *de Novo* Transformants. G3: Genes|Genomes|Genetics 6, 2829–2838.

Labib, K. (2010). How do Cdc7 and cyclin-dependent kinases trigger the initiation of chromosome replication in eukaryotic cells? Genes & Development 24, 1208–1219.

Langley, A.R., Smith, J.C., Stemple, D.L., and Harvey, S.A. (2014). New insights into the maternal to zygotic transition. Development 141, 3834–3841.

Lima-de-Faria, A., and Jaworksa, H. (1968). Late DNA Synthesis in Heterochromatin. Nature 217.

Mantiero, D., Mackenzie, A., Donaldson, A., and Zegerman, P. (2011). Limiting replication initiation factors execute the temporal programme of origin firing in budding yeast. EMBO J 30, 4805–4814.

McCune, H.J., Danielson, L.S., Alvino, G.M., Collingwood, D., Delrow, J.J., Fangman, W.L., Brewer, B.J., and Raghuraman, M.K. (2008). The Temporal Program of Chromosome Replication: Genomewide Replication in clb5 Saccharomyces cerevisiae. Genetics 180, 1833–1847.

McGuffee, S.R., Smith, D.J., and Whitehouse, I. (2013). Quantitative, Genome-Wide Analysis of Eukaryotic Replication Initiation and Termination. Molecular Cell 50, 123–135.

Melters, D.P., Bradnam, K.R., Young, H.A., Telis, N., May, M.R., Ruby, J., Sebra, R., Peluso, P., Eid, J., Rank, D., et al. (2013). Comparative analysis of tandem repeats from hundreds of species reveals unique insights into centromere evolution. Genome Biology 14, R10.

Miller, C.A., and Kowalski, D. (1993). cis-acting components in the replication origin from ribosomal DNA of Saccharomyces cerevisiae. Mol. Cell. Biol. 13, 5360–5369.

Müller, C.A., Hawkins, M., Retkute, R., Malla, S., Wilson, R., Blythe, M.J., Nakato, R., Komata, M., Shirahige, K., de Moura, A.P.S., et al. (2014). The dynamics of genome replication using deep sequencing. Nucleic Acids Res 42, e3.

Muller, M., Lucchini, R., and Sogo, J.M. (2000). Replication of Yeast rDNA Initiates Downstream of Transcriptionally Active Genes. Molecular Cell 5, 767–777.

Muramatsu, S., Hirai, K., Tak, Y.-S., Kamimura, Y., and Araki, H. (2010). CDK-dependent complex formation between replication proteins Dpb11, Sld2, Pole, and GINS in budding yeast. Genes & Development 24, 602–612.

Newlon, C.S., Lipchitz, L.R., Collins, I., Deshpande, A., Devenish, R.J., Green, R.P., Klein, H.L., Palzkill, T.G., Ren, R., Synn, S., et al. (1991). Analysis of a Circular Derivative of Saccharomyces Cerevisiae Chromosome III: A Physical Map and Identification and Location of Ars Elements. Genetics 129, 343–357.

Nishimura, K., Fukagawa, T., Takisawa, H., Kakimoto, T., and Kanemaki, M. (2009). An auxin-based degron system for the rapid depletion of proteins in nonplant cells. Nature Methods 6, 917–922.

Payen, C., Di Rienzi, S.C., Ong, G.T., Pogachar, J.L., Sanchez, J.C., Sunshine, A.B., Raghuraman, M.K., Brewer, B.J., and Dunham, M.J. (2013). The Dynamics of Diverse Segmental Amplifications in Populations of Saccharomyces cerevisiae Adapting to Strong Selection. G3 (Bethesda) 4, 399–409.

Petryk, N., Kahli, M., d’Aubenton-Carafa, Y., Jaszczyszyn, Y., Shen, Y., Silvain, M., Thermes, C., Chen, C.-L., and Hyrien, O. (2016). Replication landscape of the human genome. Nature Communications 7, 10208.

Pohl, T.J., Brewer, B.J., and Raghuraman, M.K. (2012). Functional Centromeres Determine the Activation Time of Pericentric Origins of DNA Replication in Saccharomyces cerevisiae. PLoS Genetics 8, e1002677.

Prioleau, M.-N., and MacAlpine, D.M. (2016). DNA replication origins—where do we begin? Genes Dev 30, 1683–1697.

Promega (2018). I-PpoI (Intron-Encoded Endonuclease). http://www.promega.com/products/cloning-and-dna-markers/restriction-enzymes/i_ppoi-_intron_encoded-endonuclease_/

Raghuraman, M.K., Winzeler, E.A., Collingwood, D., Hunt, S., Wodicka, L., Conway, A., Lockhart, D., Davis, R.W., Brewer, B.J., and Fangman, W.L. (2001). Replication dynamics of the yeast genome. Science 294, 115–121.

Rhind, N., and Gilbert, D.M. (2013). DNA Replication Timing. Cold Spring Harb Perspect Biol 5, a010132.

Rose, M.D. (1990). Methods in yeast genetics: a laboratory course manual (Cold Spring Harbor, N.Y.: Cold Spring Harbor Laboratory Press).

Salim, D., Bradford, W.D., Freeland, A., Cady, G., Wang, J., Pruitt, S.C., and Gerton, J.L. (2017). DNA replication stress restricts ribosomal DNA copy number. PLoS Genet 13.

Sanchez, J.C., Kwan, E.X., Pohl, T.J., Amemiya, H.M., Raghuraman, M.K., and Brewer, B.J. (2017). Defective replication initiation results in locus specific chromosome breakage and a ribosomal RNA deficiency in yeast. PLOS Genetics 13, e1007041.

Sheu, Y.-J., and Stillman, B. (2010). The Dbf4-Cdc7 kinase promotes S phase by alleviating an inhibitory activity in Mcm4. Nature 463, 113–117.

Sheu, Y.-J., Kinney, J.B., Lengronne, A., Pasero, P., and Stillman, B. (2014). Domain within the helicase subunit Mcm4 integrates multiple kinase signals to control DNA replication initiation and fork progression. Proc Natl Acad Sci U S A 111, E1899–E1908.

Shimada, K., Pasero, P., and Gasser, S.M. (2002). ORC and the intra-S-phase checkpoint: a threshold regulates Rad53p activation in S phase. Genes Dev. 16, 3236–3252.

Siefert, J.C., Georgescu, C., Wren, J.D., Koren, A., and Sansam, C.L. (2017). DNA replication timing during development anticipates transcriptional programs and parallels enhancer activation. Genome Res. 27, 1406–1416.

Siow, C.C., Nieduszynska, S.R., Muller, C.A., and Nieduszynski, C.A. (2012). OriDB, the DNA replication origin database updated and extended. Nucleic Acids Research 40, D682–D686.

Stults, D.M., Killen, M.W., Pierce, H.H., and Pierce, A.J. (2008). Genomic architecture and inheritance of human ribosomal RNA gene clusters. Genome Res. 18, 13–18.

Sun, J., Evrin, C., Samel, S., Fernández-Cid, A., Riera, A., Kawakami, H., Stillman, B., Speck, C., and Li, H. (2013). Cryo-EM structure of a helicase loading intermediate containing ORC-Cdc6-Cdt1-MCM2-7 bound to DNA. Nat Struct Mol Biol 20, 944–951.

Tanaka, S., Umemori, T., Hirai, K., Muramatsu, S., Kamimura, Y., and Araki, H. (2007). CDK-dependent phosphorylation of Sld2 and Sld3 initiates DNA replication in budding yeast. Nature 445, 328–332.

Tanaka, S., Nakato, R., Katou, Y., Shirahige, K., and Araki, H. (2011). Origin Association of Sld3, Sld7, and Cdc45 Proteins Is a Key Step for Determination of Origin-Firing Timing. Current Biology 21, 2055–2063.

Taylor, J.H. (1960). Asynchronous Duplication of Chromosomes in Cultured Cells of Chinese Hamster. J Biophys Biochem Cytol 7, 455–463.

Tercero, J.A., Longhese, M.P., and Diffley, J.F. (2003). A central role for DNA replication forks in checkpoint activation and response. Molecular Cell 11, 1323–1336.

Venema, J., and Tollervey, D. (1999). Ribosome Synthesis in Saccharomyces cerevisiae. Annual Review of Genetics 33, 261–311.

Vogelauer, M., Rubbi, L., Lucas, I., Brewer, B.J., and Grunstein, M. (2002). Histone Acetylation Regulates the Time of Replication Origin Firing. Molecular Cell 10, 1223–1233.

Willard, H.F. (1990). Centromeres of mammalian chromosomes. Trends in Genetics 6, 410–416.

Yoshida, K., Bacal, J., Desmarais, D., Padioleau, I., Tsaponina, O., Chabes, A., Pantesco, V., Dubois, E., Parrinello, H., Skrzypczak, M., et al. (2014). The Histone Deacetylases Sir2 and Rpd3 Act on Ribosomal DNA to Control the Replication Program in Budding Yeast. Molecular Cell 54, 691–697.

Zegerman, P., and Diffley, J.F.X. (2007). Phosphorylation of Sld2 and Sld3 by cyclin-dependent kinases promotes DNA replication in budding yeast. Nature 445, 281–285.

Zhang, Y., and Hunter, T. (2014). Roles of Chk1 in Cell Biology and Cancer Therapy. Int J Cancer 134.

Zhong, Y., Nellimoottil, T., Peace, J.M., Knott, S.R.V., Villwock, S.K., Yee, J.M., Jancuska, J.M., Rege, S., Tecklenburg, M., Sclafani, R.A., et al. (2013). The level of origin firing inversely affects the rate of replication fork progression. The Journal of Cell Biology 201, 373–383.

